# Comparative Mitogenomics and Molecular Phylogeny of Agriculturally Significant Tephritid Fruit Fly Pests in Bangladesh

**DOI:** 10.64898/2026.09.12.751165

**Authors:** Sadniman Rahman, Faiza Anzum Shormi

## Abstract

Tephritid fruit flies of the tribe Dacini rank among the most economically destructive agricultural pests globally, with several species of *Bactrocera* Macquart and *Zeugodacus* Hendel causing severe losses to fruit and vegetable production across South and Southeast Asia. Bangladesh harbors several economically important dacine species, including *Bactrocera dorsalis* (Hendel), *B. carambolae* (Drew & Hancock), *B. zonata* (Saunders), *B. correcta* (Bezzi) and *Zeugodacus cucurbitae* (Coquillett). Here we present a comparative mitogenomic and phylogenetic framework based on 21 unique GenBank mitogenome records: 19 dacine ingroup records and two outgroups. The ingroup includes *B. papayae, B. philippinensis* and *B. invadens*, whose synonymy with *B. dorsalis* is disputed. All records contained 13 protein-coding genes (PCGs) and two rRNA genes. Genome size in the 14 taxa of subgenus *Bactrocera* ranged from 15,273 to 15,977 bp. Whole-genome AT content ranged from 66.6% in *B. tsuneonis* to 82.2% in *Drosophila melanogaster*, and the five focal pest species clustered at 72.9–73.6%. Maximum-likelihood analysis of a 13,600-site alignment (13 PCGs plus 12S and 16S rRNA; GTR+FO; IQ-TREE v1.6.11; 1,000 ultrafast bootstrap replicates, UFBoot) recovered the *B. dorsalis* complex (UFBoot = 100), within which pairwise distances were only 0.64–1.33%, and a *B. correcta* + *B. zonata* pair (UFBoot = 100). Nine taxa showed compositional heterogeneity and third codon positions were highly variable, so deeper nodes should be interpreted cautiously. As no Bangladeshi specimens were sequenced, this study is a curated reference baseline for molecular identification and pest surveillance, not a population-genomic analysis.

## 1. Introduction

Fruit flies of the tribe Dacini (Diptera: Tephritidae) include some of the world’s most economically destructive agricultural pests. The tribe comprises 932 recognized species, about 10% of which are pests of commercial fruits and vegetables whose presence triggers quarantines and trade restrictions (Doorenweerd et al., 2018). Most of these pests belong to *Bactrocera* Macquart, 1835, which in its broad sense contains more than 650 described species (Vargas et al., 2015), or to *Zeugodacus* Hendel, 1927, which was long treated as a subgenus of *Bactrocera* but is now recognized as a separate genus on molecular and morphological grounds (Krosch et al., 2012; Virgilio et al., 2015; San Jose et al., 2018a; Doorenweerd et al., 2018). Their larvae damage a wide range of fruit and vegetable crops, and several species have spread far beyond their native ranges (Clarke et al., 2005; Vargas et al., 2015).

Bangladesh supports a diverse Dacini fauna. Six years of surveys (2013–2018) raised the number of tephritid species recorded from the country from seven to 37 and documented the first records of the highly invasive *B. carambolae* Drew & Hancock, 1994 (Leblanc et al., 2019). Five dacine species are considered here as focal pests: the polyphagous fruit pests *B. dorsalis* Hendel, 1912, *B. zonata* Saunders, 1842, *B. correcta* Bezzi, 1916 and *B. carambolae*, and the cucurbit pest *Zeugodacus cucurbitae* Coquillett, 1899 (Leblanc et al., 2019). *B. dorsalis* and *B. carambolae* belong to the *B. dorsalis* species complex, a group of polyphagous pests of worldwide quarantine concern (Clarke et al., 2005). Species limits in this complex have been difficult to resolve: *B. papayae, B. philippinensis* and *B. invadens* were synonymized with *B. dorsalis* by Schutze et al. (2015a, 2015b), a treatment that has since been contested (Schutze et al., 2017), and mitochondrial COI sequences cannot separate *B. dorsalis* from *B. carambolae*, so that nuclear markers are needed to identify them (San Jose et al., 2018b; Leblanc et al., 2019).

Complete mitochondrial genomes (mitogenomes) are widely used in insect systematics and pest identification because they are compact, uniparentally inherited and contain a standard set of 13 protein-coding genes (PCGs), 22 transfer RNA genes and two ribosomal RNA genes, together with a non-coding control region (Boore, 1999; Cameron, 2014). In Dacini, mitogenomes have been described for a growing number of species, including *B. minax* (Zhang et al., 2014), *B. arecae* (Yong et al., 2015), *B. latifrons, B. melastomatos* and *B. umbrosa* (Yong et al., 2016), *B. invadens* (Zhang et al., 2016), *B. carambolae* (Drosopoulou et al., 2019), *B. tuberculata* (Wang et al., 2020) and *B. cinnabaria* and *B. propinqua* (Yong et al., 2024), and a recent survey of 82 *Bactrocera* mitogenomes representing 16 species has documented their structure and diagnostic potential (Castellanos et al., 2025). To our knowledge, however, no mitogenomic comparison has been framed around the pest assemblage of Bangladesh, and the amount of mitogenome divergence among close relatives in the *B. dorsalis* complex, relative to that among the other Bangladeshi pests, has not been quantified.

Here we assemble a curated 21-taxon dataset from GenBank in which eight dacine species recorded from Bangladesh are represented together with comparators and two outgroups. We (i) quality-control the records and describe genome size, nucleotide composition, strand asymmetry and control-region annotation; (ii) infer a maximum-likelihood phylogeny from a concatenated alignment of the 13 PCGs and the two rRNA genes; and (iii) quantify pairwise divergence among the Bangladeshi pests and the *B. dorsalis* complex and assess the compositional and saturation properties of the alignment. These results provide a comparative reference framework, not a population-genomic analysis.

## 2. Materials and Methods

### 2.1 Dataset and Sequence Quality Control

A GenBank file with 25 records was parsed with Biopython (Cock et al., 2009). After removing duplicates (NC_009772.1 occurred four times and NC_009771.1 twice), 21 unique accession.version identifiers remained (Table 1). The dataset comprises 14 taxa assigned to *Bactrocera* sensu stricto, *Z. cucurbitae*, four further dacine comparators (*B. minax, B. tsuneonis, B. cinnabaria* and *B. oleae*, representing the groups *Tetradacus, Bulladacus* and *Daculus*) and two non-dacine outgroups, *Ceratitis capitata* and *D. melanogaster*. Group labels follow the traditional subgeneric assignments and are used descriptively, because several traditional subgenera are not monophyletic (San Jose et al., 2018a). Eight dacine species in the dataset have been recorded from Bangladesh (Leblanc et al., 2019): the five focal pests (Table 2) and *B. tuberculata* Bezzi, *B. propinqua* Hardy & Adachi and *B. latifrons* Hendel; the remaining taxa serve as phylogenetic comparators. *B. papayae, B. philippinensis* and *B. invadens* are retained under their GenBank names although they were synonymized with *B. dorsalis* by Schutze et al. (2015a, 2015b), a treatment that remains contested (Schutze et al., 2017). All 21 records contained 13 CDS, 22 tRNA and two rRNA features. MT196006 (*B. tuberculata*) is labelled “partial genome” in GenBank but is described as a complete 15,273-bp mitogenome in its original publication (Wang et al., 2020); it was retained.

**Table 1.** Mitogenome features of the 21 records analysed. Length, AT content and strand skews were computed from the GenBank sequences (skews for the strand as deposited). CR, annotated control-region (D-loop) length; NA, no annotation in the supplied record. Focal pest, the five species detailed in Table 2; Recorded in Bangladesh, other dataset species recorded by Leblanc et al. (2019). B., Bactrocera; Z., Zeugodacus; C., Ceratitis; D., Drosophila. B. papayae, B. philippinensis and B. invadens were synonymized with B. dorsalis by Schutze et al. (2015a, 2015b); this treatment is contested (Schutze et al., 2017).

| **Taxon** | **Accession** | **Length (bp)** | **AT (%)** | **AT skew** | **GC skew** | **CR (bp)** | **Group** | **Status** |
|---|---|---|---|---|---|---|---|---|
| *B. dorsalis* | NC_008748.1 | 15,915 | 73.58 | 0.068 | –0.228 | 949 | *Bactrocera* s.s. | Focal pest |
| *B. carambolae* | NC_009772.1 | 15,915 | 73.55 | 0.066 | –0.224 | 950 | *Bactrocera* s.s. | Focal pest |
| *B. papayae* | NC_009770.1 | 15,915 | 73.52 | 0.066 | –0.226 | 950 | *Bactrocera* s.s. | Comparator |
| *B. philippinensis* | NC_009771.1 | 15,915 | 73.63 | 0.066 | –0.224 | 949 | *Bactrocera* s.s. | Comparator |
| *B. correcta* | NC_018787.1 | 15,936 | 73.17 | 0.063 | –0.222 | 949 | *Bactrocera* s.s. | Focal pest |
| *B. zonata* | NC_027725.1 | 15,935 | 73.34 | 0.065 | –0.223 | 950 | *Bactrocera* s.s. | Focal pest |
| *B. tryoni* | NC_014611.1 | 15,925 | 72.42 | 0.070 | –0.227 | NA | *Bactrocera* s.s. | Comparator |
| *B. latifrons* | NC_029466.1 | 15,977 | 71.11 | 0.088 | –0.266 | 953 | *Bactrocera* s.s. | Recorded in Bangladesh |
| *B. melastomatos* | NC_029467.1 | 15,954 | 73.79 | 0.073 | –0.251 | 953 | *Bactrocera* s.s. | Comparator |
| *B. umbrosa* | NC_029468.1 | 15,898 | 70.49 | 0.084 | –0.244 | 944 | *Bactrocera* s.s. | Comparator |
| *B. arecae* | NC_028327.1 | 15,900 | 72.26 | 0.081 | –0.259 | 952 | *Bactrocera* s.s. | Comparator |
| *B. invadens* | NC_031388.1 | 15,911 | 73.63 | 0.067 | –0.229 | 949 | *Bactrocera* s.s. | Comparator |
| *B. tuberculata* | MT196006.1 | 15,273 | 72.26 | 0.063 | –0.215 | 251 | *Bactrocera* s.s. | Recorded in Bangladesh |
| *B. propinqua* | OR085850.1 | 15,927 | 74.05 | 0.069 | –0.226 | 947 | *Bactrocera* s.s. | Recorded in Bangladesh |
| *Z. cucurbitae* | NC_016056.1 | 15,825 | 72.89 | 0.047 | –0.213 | 946 | *Zeugodacus* | Focal pest |
| *B. minax* | NC_014402.1 | 16,043 | 67.28 | 0.131 | –0.316 | NA | *Tetradacus* | Comparator |
| *B. tsuneonis* | NC_038164.1 | 15,865 | 66.57 | 0.150 | –0.340 | 949 | *Tetradacus* | Comparator |
| *B. cinnabaria* | OR085849.1 | 15,225 | 71.16 | 0.085 | –0.250 | 358 | *Bulladacus* | Comparator |
| *B. oleae* | NC_005333.1 | 15,815 | 72.63 | 0.088 | –0.280 | 949 | *Daculus* | Comparator |
| *C. capitata* | NC_000857.1 | 15,980 | 77.48 | 0.021 | –0.185 | NA | Outgroup | Outgroup |
| *D. melanogaster* | NC_024511.2 | 19,524 | 82.22 | 0.016 | –0.147 | 4,608 | Outgroup | Outgroup |

**Table 2.**
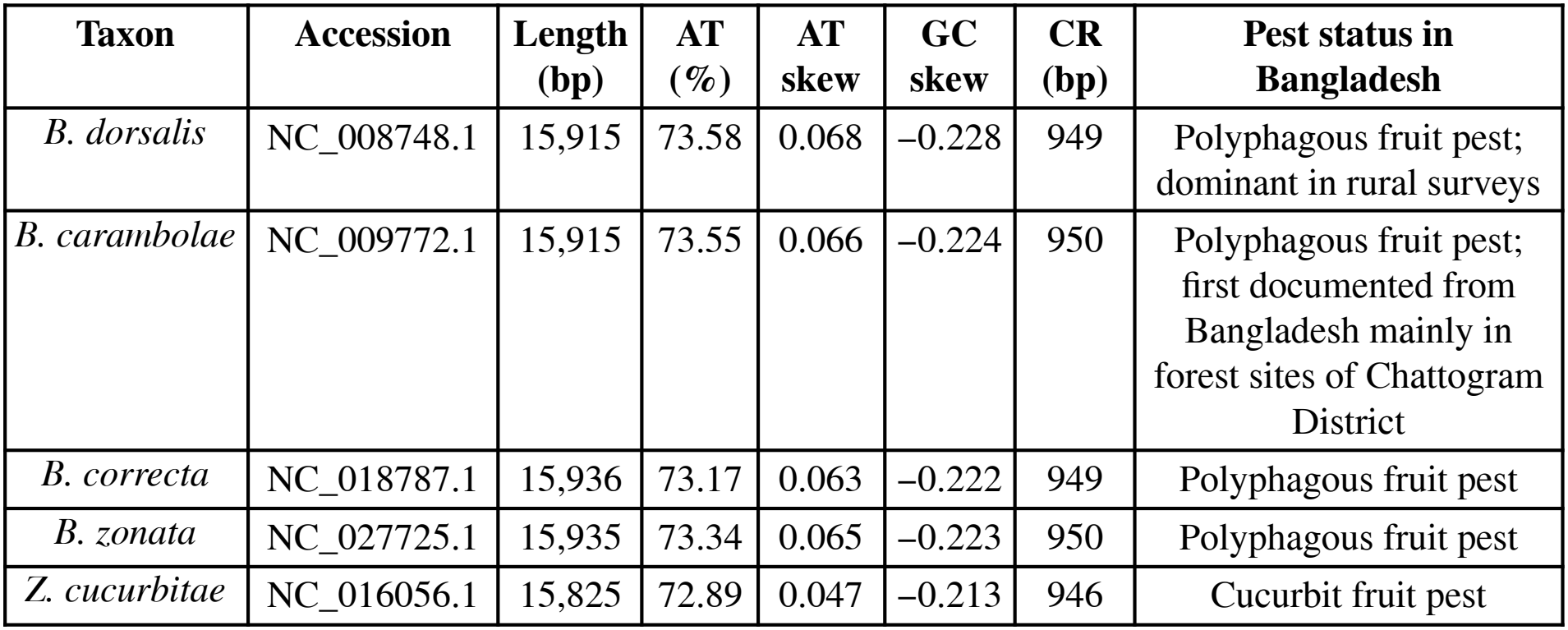
The five focal pest species. Values as in Table 1. Pest status follows Leblanc et al. (2019).

### 2.2 Genome Statistics

Genome length, AT and GC content, AT skew [(A − T)/(A + T)] and GC skew [(G − C)/(G + C)] were computed from the nucleotide sequences as deposited in GenBank, that is, for the strand of the submitted sequence. Control-region length is the length of the annotated D-loop or misc_feature record where one was present. Feature counts (CDS, tRNA, rRNA) were tallied per record.

### 2.3 Gene Extraction and Alignment

Each of the 13 PCGs was extracted from the CDS annotations (respecting strand), translated with the invertebrate mitochondrial genetic code (NCBI translation table 5; Elzanowski & Ostell, 2024) and stripped of its terminal stop codon. Each protein was aligned to the corresponding *B. dorsalis* (NC_008748.1) protein with a global pairwise alignment (Biopython PairwiseAligner; match +2, mismatch −1, gap-open −8, gap-extend −0.5). The pairwise alignments were merged into a reference-anchored (“star”) alignment that retains insertions relative to the reference and were back-translated to the original codons, so that all gaps in the PCG block span whole codons. The 12S and 16S rRNA genes were aligned in the same way at the nucleotide level. Blocks were concatenated in the order *nad2, cox1, cox2, atp8, atp6, cox3, nad3, nad5, nad4, nad4l, nad6, cytb, nad1*, 16S, 12S, giving 13,600 columns (PCG block 11,283 columns = 3,761 codons; 16S 1,451; 12S 866; Table 3). No trimming or masking was applied; gaps and ambiguity codes make up 2.38% of the matrix on average. Because this is a reference-anchored alignment and not an iterative multiple-sequence alignment, alignment uncertainty in divergent regions and taxa, especially the outgroups, cannot be excluded (Section 4).

**Table 3.** Summary of the alignment and maximum-likelihood analysis.

| **Parameter** | **Value** |
|---|---|
| Taxa | 21 (19 dacine ingroup records, 2 outgroups) |
| Alignment method | Reference-anchored (star) codon-aware alignment to B. dorsalis NC_008748.1 (Biopython PairwiseAligner; protein-guided for PCGs, nucleotide for rRNAs); no trimming |
| Alignment length | 13,600 columns |
| PCG block | 11,283 columns (3,761 codons; 13 genes) |
| 16S rRNA block | 1,451 columns |
| 12S rRNA block | 866 columns |
| Gaps and ambiguity | 2.38% of the matrix (mean) |
| Constant sites | 7,232 (53.2%) |
| Parsimony-informative sites | 4,027 (29.6%) |
| Variable, not parsimony-informative | 2,341 (17.2%) |
| Distinct site patterns | 4,033 |
| Model | GTR+FO (general time-reversible, empirical base frequencies); unpartitioned |
| Software | IQ-TREE v1.6.11 (Nguyen et al., 2015) |
| Branch support | 1,000 ultrafast bootstrap replicates (Hoang et al., 2018) |
| Command | -st DNA -m GTR+FO -bb 1000 -pers 0.5 -numstop 100 |
| Log-likelihood of best tree | –101,310.401 |
| Total tree length | 1.520 substitutions per site |
| Search convergence | 102 iterations; UFBoot correlation coefficient 0.998 |
| Composition chi-square test | 9 of 21 sequences failed (P < 0.05): *B. latifrons*, *B. umbrosa*, *B. tuberculata*, *B. propinqua*, *B. minax*, *B. tsuneonis*, *B. cinnabaria*, *C. capitata*, *D. melanogaster* |
| Display root | *D. melanogaster* NC_024511.2 |

### 2.4 Phylogenetic Analysis

The 21-taxon matrix was analyzed with IQ-TREE v1.6.11 (Nguyen et al., 2015) under the GTR+FO model (general time-reversible, empirical base frequencies) with 1,000 ultrafast bootstrap replicates (UFBoot; Hoang et al., 2018) (command: -st DNA -m GTR+FO -bb 1000 -pers 0.5 -numstop 100). A single homogeneous model was applied to the whole matrix; no partitioning and no automated model selection were used. The search converged after 102 iterations (UFBoot correlation coefficient 0.998; Table 3). IQ-TREE’s composition chi-square test flagged nine sequences (*B. latifrons, B. umbrosa, B. tuberculata, B. propinqua, B. minax, B. tsuneonis, B. cinnabaria, C. capitata* and *D. melanogaster*; P < 0.05), so deeper relationships involving these taxa are interpreted cautiously. Support values are properties of the splits of the unrooted tree; for display, the tree was rooted on *D. melanogaster* and each clade was assigned the support of the corresponding split.

### 2.5 Pairwise Divergence and Alignment Properties

Uncorrected pairwise p-distances were computed from the alignment with pairwise deletion of gaps, for the whole matrix and separately for codon positions 1 + 2 and for position 3 of the PCG block (codon position follows from the column index because all gaps in the PCG block are in frame). Sites were classified per region as constant, variable but not parsimony-informative, or parsimony-informative, with gaps treated as missing; the overall counts reproduce those reported by IQ-TREE (7,232, 2,341 and 4,027 sites). The full distance matrix is provided as Supplementary Table S1.

## 3. Results

### 3.1 Dataset and Genome Size

After duplicate removal the dataset comprised 21 records, each with 13 annotated CDS, 22 tRNA and two rRNA features (Table 1). Mitogenome length ranged from 15,225 bp (*B. cinnabaria*) to 19,524 bp (*D. melanogaster*), and from 15,273 bp (*B. tuberculata*) to 15,977 bp (*B. latifrons*) among the 14 *Bactrocera* s.s. taxa (mean ± SD, 15,878.3 ± 175.5 bp; Fig. 1A). The five focal species spanned 15,825–15,936 bp (Table 2). The shortest records were *B. cinnabaria* (15,225 bp) and *B. tuberculata* (15,273 bp), the latter labelled as a partial genome in GenBank.

**Fig. 1.**
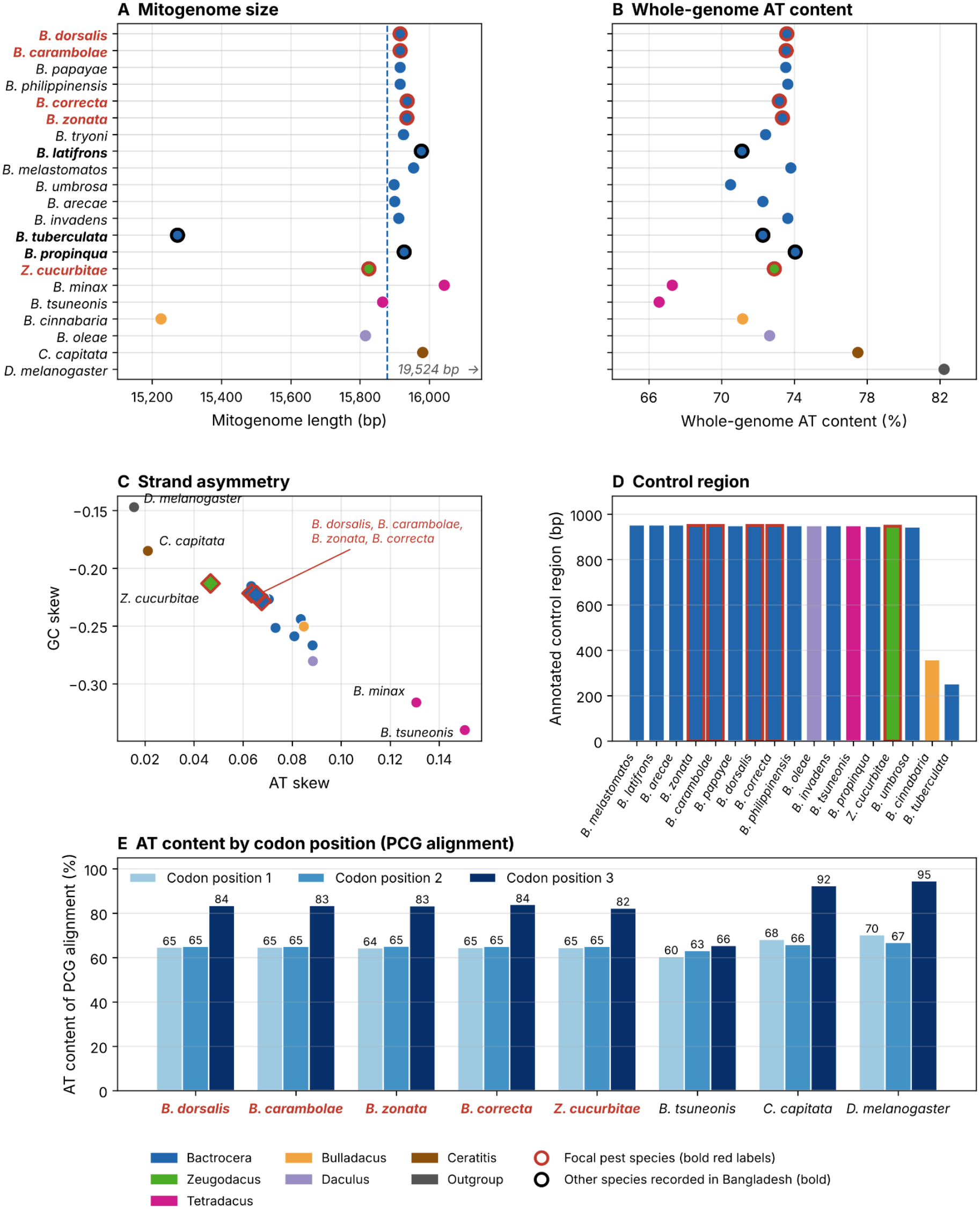
Genome features of the 21 records, computed from the GenBank sequences and the concatenated alignment. (A) Mitogenome length (bp); the dashed line is the mean of the 14 Bactrocera s.s. taxa (15,878 bp) and D. melanogaster (19,524 bp) lie beyond the axis. (B) Whole-genome AT content (%). (C) AT skew versus GC skew, for the strand as deposited; red diamonds are the five focal species, whose four Bactrocera members overlap. (D) Annotated control-region (D-loop) length for the 17 records with an annotation, excluding D. melanogaster (4,608 bp); B. tryoni, B. minax and C. capitata have no annotated control region. (E) AT content (%) at codon positions 1, 2 and 3 of the PCG block of the alignment for eight selected taxa. Colours indicate group. Red outlines and bold red labels mark the five focal pest species; black outlines and bold labels mark the other dataset species recorded from Bangladesh (B. tuberculata, B. propinqua, B. latifrons).

### 3.2 Nucleotide Composition and Strand Asymmetry

Whole-genome AT content was 70.49–74.05% among the 14 *Bactrocera* s.s. taxa (mean 72.91%), 72.89% in *Z. cucurbitae*, 71.16% in *B. cinnabaria*, 72.63% in *B. oleae*, 77.48% in *C. capitata* and 82.22% in *D. melanogaster* (Fig. 1B). The five focal species were tightly clustered (72.89–73.58%; Table 2). The two *Tetradacus* records were markedly more GC-rich than any other ingroup taxon (*B. minax*, 67.28%; *B. tsuneonis*, 66.57%), a pattern that coincides with their failure of IQ-TREE’s composition test (Section 2.4). Every record showed a positive AT skew (0.016 to 0.150) and a negative GC skew (−0.147 to −0.340) for the strand as deposited (Fig. 1C). The four focal *Bactrocera* species were nearly indistinguishable (AT skew 0.063–0.068; GC skew −0.222 to −0.228), whereas *Z. cucurbitae* differed (AT skew 0.047; GC skew −0.213). Strand asymmetry of this sign is typical of insect mitogenomes and has been attributed to strand-biased replication and transcription-associated mutation, although its basis and its reversals across lineages remain debated (Reyes et al., 1998; Hassanin et al., 2005).

### 3.3 Control Region Annotation

An annotated control region (D-loop or misc_feature record) was present in 18 of 21 records; *B. tryoni, B. minax* and *C. capitata* lacked one. Excluding *D. melanogaster* (4,608 bp), the 17 annotated lengths were 944–953 bp in 15 records, including all five focal species (946–950 bp), and shorter in *B. tuberculata* (251 bp) and *B. cinnabaria* (358 bp) (Fig. 1D). The near identity of annotated lengths among distantly related taxa suggests that annotation conventions contribute as much as biological conservation, so these values should not be used as diagnostic characters; the short values in *B. tuberculata* and *B. cinnabaria* may reflect incomplete control regions.

### 3.4 Gene Content and Compositional Structure of the Alignment

All 21 records contained the canonical complement of 13 PCGs, 22 tRNA genes and two rRNA genes. The supplied GenBank annotations place *atp6, atp8, cox1, cox2, cox3, cytb, nad2, nad3* and *nad6* on the majority (J) strand and *nad1, nad4, nad4l* and *nad5*, with the two rRNAs, on the minority (N) strand, the arrangement typical of insect mitogenomes. In the concatenated alignment, AT content of the PCG block was 70.6–71.2% in the focal species, rising from 64.3–65.1% at codon positions 1 and 2 to 82.3–83.9% at position 3 (Fig. 1E); the rRNA blocks were more AT-rich than the PCG block (16S, 79.2–79.8%; 12S, 74.8–75.1%). The *Tetradacus* record *B. tsuneonis* was strikingly different, with only 65.5% AT at third positions (60.5% and 63.2% at positions 1 and 2), whereas the outgroups showed the strongest third-position bias (*C. capitata*, 92.4%; *D. melanogaster*, 94.5%). These contrasts quantify the compositional heterogeneity detected by IQ-TREE.

### 3.5 Phylogenetic Relationships

The ML tree (GTR+FO; log-likelihood −101,310.40; tree length 1.520; Table 3) is shown in Fig. 2 with *D. melanogaster* as the root; each number is the UFBoot support of the corresponding clade. (i) Excluding *C. capitata*, the 19 ingroup taxa formed a clade (UFBoot = 100), within which *Z. cucurbitae* was sister to the remaining 18 *Bactrocera*-group taxa (96). (ii) The 14 *Bactrocera* s.s. taxa were monophyletic (100). The other four dacine comparators formed a separate clade (86), within which *B. minax* and *B. tsuneonis* were sister taxa (100) and *B. cinnabaria* joined them with weaker support (69). (iii) Within *Bactrocera* s.s., *B. latifrons* and *B. umbrosa* formed a pair (71) sister to the remaining 12 taxa (62); *B. propinqua* was sister to the remaining 11 (80); *B. tryoni* and *B. arecae* formed a pair (69), and the nine-taxon group that excludes them (67) comprised *B. correcta* and *B. zonata* (100), which together with *B. melastomatos* (56) were sister to *B. tuberculata* plus the *B. dorsalis* complex (84). (iv) The five names of the *B. dorsalis* complex (*B. dorsalis, B. invadens, B. papayae, B. philippinensis* and *B. carambolae*) formed a clade (100). Within it, *B. carambolae* was sister to a four-name group (79) that comprised the pairs (*B. dorsalis, B. invadens*) (85) and (*B. papayae, B. philippinensis*) (85). The nodes within the complex are too close together to label on the tree and are given in the box in Fig. 2. Relationships deep in the *Bactrocera* s.s. clade were weakly to moderately supported (56–84).

**Fig. 2.**
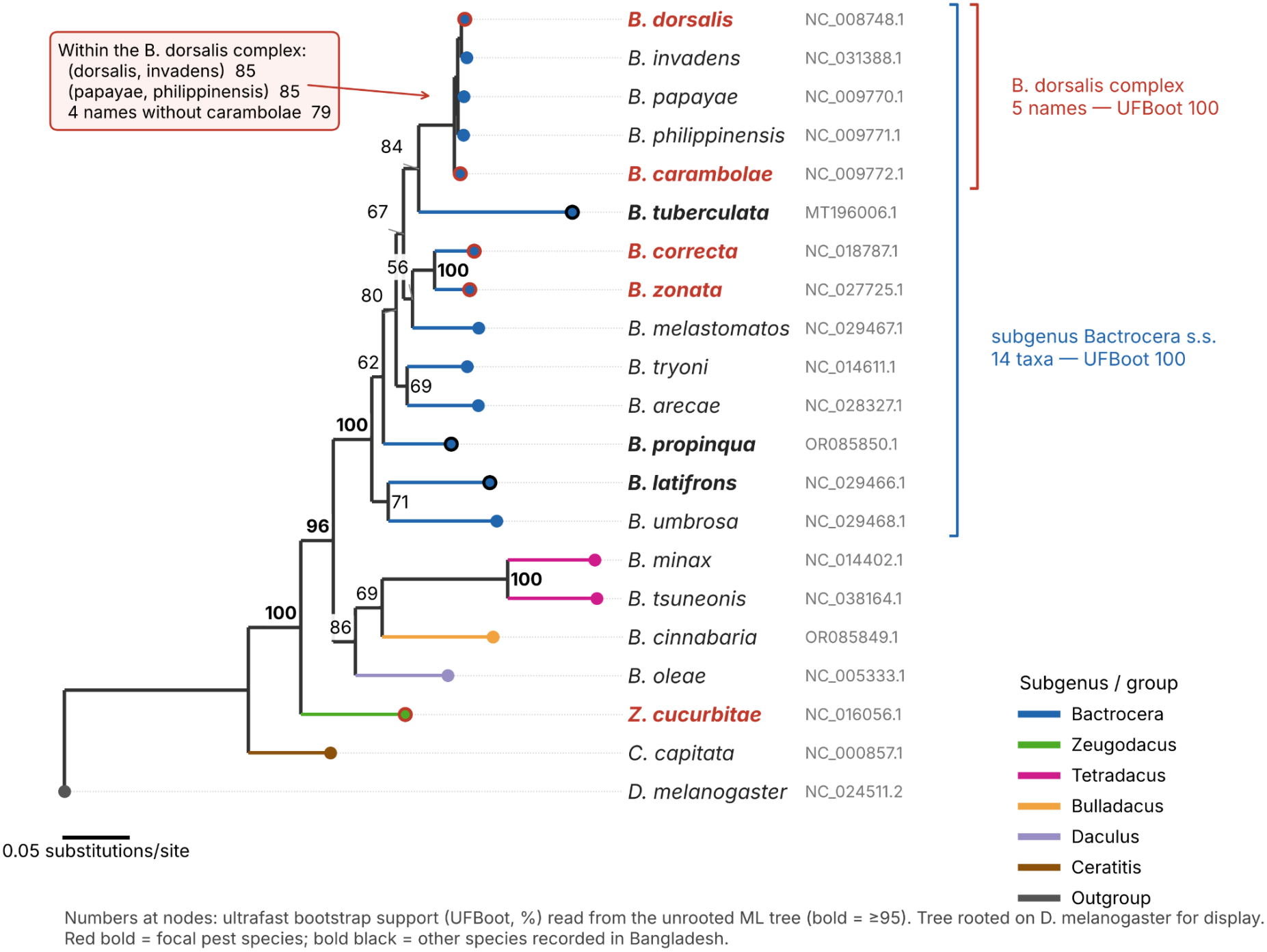
Maximum-likelihood phylogeny of 21 mitogenomes inferred from 13,600 aligned sites (13 PCGs, 12S and 16S rRNA) under GTR+FO in IQ-TREE v1.6.11 (log-likelihood −101,310.40; tree length 1.520 substitutions per site). Numbers at nodes are ultrafast bootstrap supports (UFBoot, %; 1,000 replicates; bold ≥ 95) read from the splits of the unrooted tree and assigned to the corresponding clades; supports for the nodes inside the B. dorsalis complex, which are too close together to label, are given in the box. The tree is drawn rooted on D. melanogaster for display only. Red bold labels, focal pest species; bold black labels, other species recorded from Bangladesh. Nine taxa failed IQ-TREE’s composition test (P < 0.05), so deeper nodes should be interpreted cautiously.

### 3.6 Pairwise Divergence

Among the five names of the *B. dorsalis* complex, p-distances ranged from 0.64% (*B. dorsalis*–*B. invadens*) to 1.33% (*B. carambolae*–*B. invadens*); *B. carambolae* differed from *B. dorsalis* by 1.24%, from *B. papayae* by 1.15% and from *B. philippinensis* by 1.09%, and the four names other than *B. carambolae* differed from each other by 0.64–1.16% (Fig. 3A). In contrast, *B. correcta* and *B. zonata* differed from each other by 5.54% and from members of the *B. dorsalis* complex by 7.46–7.61%, and *Z. cucurbitae* differed from all other focal species by 13.59–13.81% (Fig. 3A). Distances from *B. dorsalis* to the other taxa increased from 0.6% (*B. invadens*) to 19.0% (*D. melanogaster*); the other *Bactrocera* s.s. taxa were 7.6–13.2% distant, and *Z. cucurbitae, C. capitata, B. minax* and *B. tsuneonis* were 13.7%, 14.4%, 17.6% and 18.7% distant, respectively (Fig. 3B).

**Fig. 3.**
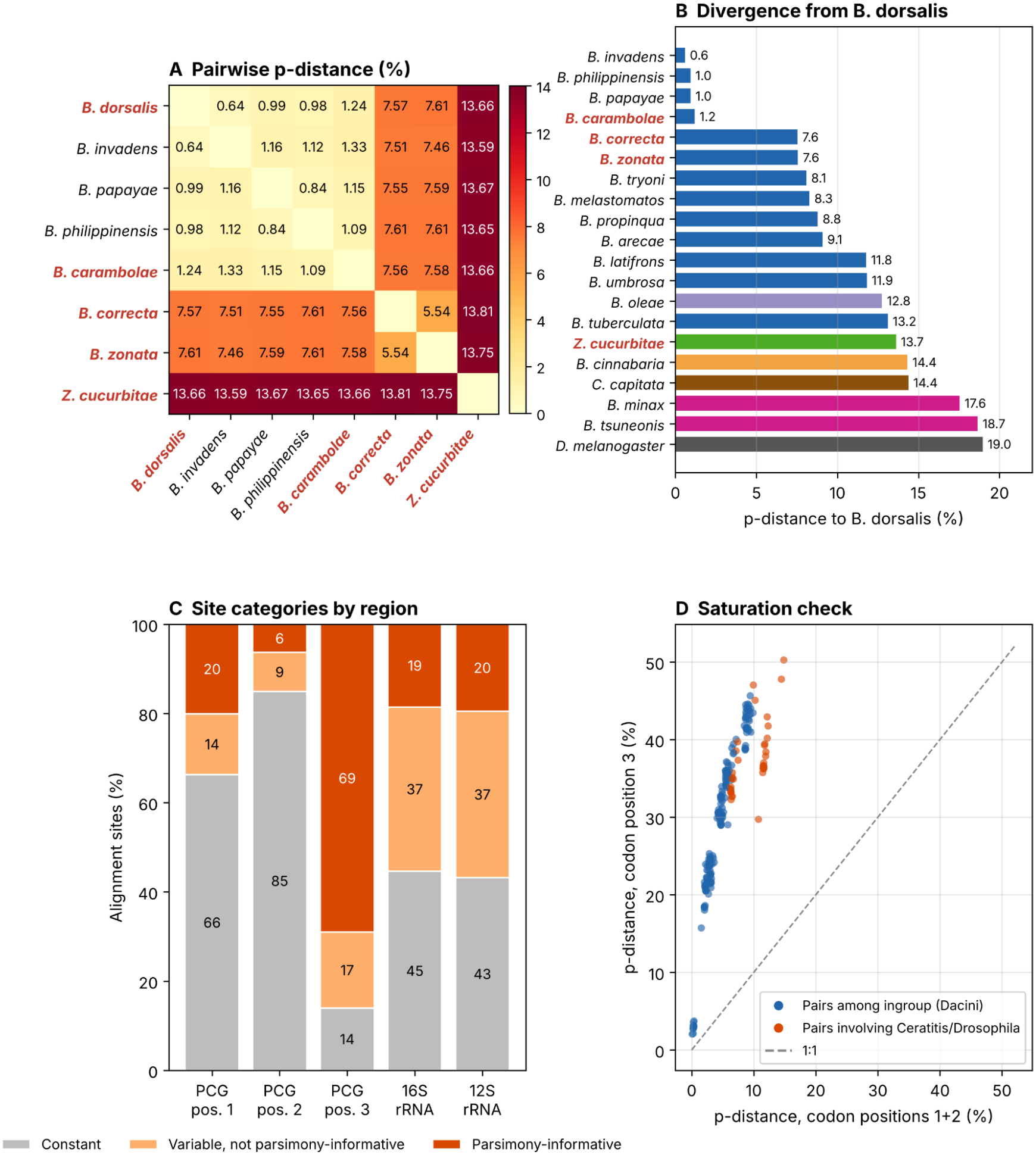
Divergence and alignment properties. (A) Pairwise uncorrected p-distance (%) among the five names of the B. dorsalis complex and the other focal pest species. (B) p-distance (%) from B. dorsalis to every other taxon, coloured by group. (C) Proportion of constant, variable (not parsimony-informative) and parsimony-informative sites by alignment region (PCG codon positions 1–3, 3,761 sites each; 16S, 1,451; 12S, 866); gaps treated as missing. (D) p-distance at codon position 3 versus positions 1 + 2 for all 210 taxon pairs; blue, pairs among the ingroup; orange, pairs involving C. capitata or D. melanogaster; dashed line, 1:1. The full distance matrix is given in Supplementary Table S1.

### 3.7 Alignment Properties and Saturation

Of the 13,600 alignment sites, 7,232 (53.2%) were constant, 2,341 (17.2%) were variable but not parsimony-informative and 4,027 (29.6%) were parsimony-informative. Variation was concentrated at third codon positions, 69.0% of which were parsimony-informative and only 14.0% constant, compared with 20.1% and 66.3% at first positions, 6.3% and 84.9% at second positions, 18.5% and 44.6% in 16S and 19.5% and 43.2% in 12S (Fig. 3C). Third-position p-distances were four to five times larger than those at positions 1 + 2 for closely related taxa and reached 45.7% among ingroup pairs, against a maximum of 9.9% at positions 1 + 2 (Fig. 3D). Pairs involving the outgroups did not have larger third-position distances (for example 36.7% for *D. melanogaster* versus *B. dorsalis*, against 11.6% at positions 1 + 2), which indicates that third positions are saturated or compositionally biased at deeper divergences.

### 3.7 Alignment Properties and Saturation

Of the 13,600 alignment sites, 7,232 (53.2%) were constant, 2,341 (17.2%) were variable but not parsimony-informative and 4,027 (29.6%) were parsimony-informative. Variation was concentrated at third codon positions, 69.0% of which were parsimony-informative and only 14.0% constant, compared with 20.1% and 66.3% at first positions, 6.3% and 84.9% at second positions, 18.5% and 44.6% in 16S and 19.5% and 43.2% in 12S (Fig. 3C). Third-position p-distances were four to five times larger than those at positions 1 + 2 for closely related taxa and reached 45.7% among ingroup pairs, against a maximum of 9.9% at positions 1 + 2 (Fig. 3D). Pairs involving the outgroups did not have larger third-position distances (for example 36.7% for *D. melanogaster* versus *B. dorsalis*, against 11.6% at positions 1 + 2), which indicates that third positions are saturated or compositionally biased at deeper divergences.

## 4. Discussion

This study provides a sequence-verified comparative framework for the dacine pests of Bangladesh. All values for genome length, nucleotide composition and strand skew were computed directly from the GenBank sequences, and the phylogenetic analysis used an explicit, reproducible pipeline. The principal finding for surveillance is how little mitogenome divergence separates the focal *B. dorsalis* from *B. carambolae* (1.24% over 13,600 aligned sites), a value comparable to those separating *B. dorsalis* from *B. papayae* (0.99%) and *B. philippinensis* (0.98%) and only about twice that between *B. dorsalis* and *B. invadens* (0.64%). By contrast, the other focal pest pair, *B. correcta* and *B. zonata*, differ by 5.54%, and these species differ from the *B. dorsalis* complex by 7.5–7.6% (Fig. 3A).

This small divergence agrees with the long-standing difficulty of separating *B. dorsalis* and *B. carambolae* with mitochondrial markers. COI barcodes do not distinguish them, and their identification has required nuclear sequences (San Jose et al., 2018b; Leblanc et al., 2019); the mitochondrial haplotypes of the two species have been described as genetically intertwined (San Jose et al., 2018b). The recognition of *B. carambolae* as a separate species rests on nuclear, morphological, behavioural and chemical evidence (Schutze et al., 2015a, 2015b). In our tree *B. carambolae* is sister to the other four names (UFBoot = 79), a moderately supported result that is consistent with, but cannot by itself establish, its distinctness. That the whole mitogenome retains some signal that separates *B. carambolae* suggests that mitogenome-scale data may offer more resolution than COI alone, but this must be tested with population-level samples, because a single reference genome per name cannot reveal haplotype sharing. The low divergence among *B. dorsalis, B. papayae, B. philippinensis* and *B. invadens* (0.64–1.16%) is compatible with their synonymy (Schutze et al., 2015a, 2015b), although mitochondrial similarity alone cannot settle a taxonomic question that remains contested (Schutze et al., 2017). For quarantine in Bangladesh, where *B. carambolae* was documented only recently (Leblanc et al., 2019), the practical implication is that mitochondrial similarity should not be relied on to rule out *B. carambolae*.

Beyond the *B. dorsalis* complex, the tree is consistent with the current classification of the tribe. Rooted on *D. melanogaster*, the ingroup excluding *C. capitata* was monophyletic (UFBoot = 100), *Z. cucurbitae* was sister to the remaining *Bactrocera*-group taxa (96) and the 14 *Bactrocera* s.s. taxa were monophyletic (100), in line with the recognition of *Zeugodacus* as a lineage distinct from *Bactrocera* (Krosch et al., 2012; Virgilio et al., 2015; San Jose et al., 2018a). The maximal support for *B. correcta* + *B. zonata* and for *B. minax* + *B. tsuneonis* adds mitogenomic support for these pairings. *B. tuberculata*, a polyphagous fruit pest recorded in Bangladesh (Leblanc et al., 2019), was recovered as sister to the *B. dorsalis* complex (84). However, it sits on a long branch (13.2% from *B. dorsalis*), failed the composition test and has a record labelled as a partial genome, so this placement is tentative.

Several features of the data call for caution. Nine of 21 sequences failed the composition test, and the Tetradacus records are conspicuously GC-rich, with third-position AT of only 65.5% in *B. tsuneonis* against 82–84% in the focal species (Fig. 1E). Third-position distances reached 45.7% and were not larger for the outgroup pairs than for some ingroup pairs (Fig. 3D), which indicates saturation. Compositional heterogeneity and saturation can produce artefactual groupings and inflate support in standard substitution models (Hassanin et al., 2005; Philippe et al., 2011). We used a single homogeneous GTR+FO model without partitioning, so nodes deep in the tree (UFBoot 56–86) and the placement of taxa with atypical composition should be considered provisional. Sensitivity analyses that exclude or RY-code third positions, partition by codon position and region, or use site-heterogeneous mixture models (Crotty et al., 2020) are the obvious next step. The rooting on a single, distant and compositionally extreme outgroup (95% AT at third positions) also limits what can be concluded about basal relationships.

The alignment is a further limitation. It was built by reference-anchored pairwise alignment to *B. dorsalis* without trimming, which is adequate for the closely related focal taxa (p-distances of 0.6–1.3% within the *B. dorsalis* complex) but may misalign divergent regions of the more distant taxa. The p-distances and site statistics for divergent pairs should therefore be read as approximate, and the analysis should be repeated with an iterative codon-aware aligner and alignment trimming. Annotated control-region lengths were nearly invariant among distantly related taxa (Section 3.3) and probably reflect annotation conventions; they are not informative for identification.

Finally, the dataset is restricted to GenBank reference genomes. Eight of the species analysed have been recorded from Bangladesh, but other important pests there, notably *Z. tau* (the third most abundant pest in rural collections in the survey of Leblanc et al., 2019), are not represented, and no Bangladeshi specimens were sequenced. The origin of *B. carambolae* in Bangladesh, whether a recent introduction or the western edge of its natural range, remains unresolved (Leblanc et al., 2019), and this study cannot address it. Complete mitogenomes of *B. dorsalis* and *B. carambolae* from replicated Bangladeshi and Myanmar populations, combined with nuclear markers, would allow haplotype sharing, population structure and invasion pathways to be assessed directly, and would show whether the low divergence reported here is typical of the two species across their ranges.

## 5. Conclusions

We provide a sequence-verified comparative reference for dacine fruit-fly mitogenomes relevant to Bangladesh, based on 21 GenBank records. All records carry the canonical insect mitochondrial gene set, and genome size, composition and strand skews are reported from the sequences themselves. Pairwise mitogenome divergence among the five names of the *B. dorsalis* complex is only 0.64–1.33%, much lower than the 5.5–7.6% separating *B. correcta, B. zonata* and the complex, which shows that mitochondrial similarity should not be used to rule out *B. carambolae*. The ML tree recovers the complex (UFBoot = 100) and a *B. correcta* + *B. zonata* pair (100), whereas deeper nodes are weakly to moderately supported and affected by compositional heterogeneity and third-position saturation. Replicated sampling of Bangladeshi populations, iterative alignment and site-heterogeneous models are needed before population-level or deep-phylogenetic conclusions can be drawn.

## Author Contributions

S.R.: Conceptualization, Methodology, Data Curation, Writing – Original Draft, Project Administration. F.A.S.: Formal Analysis, Visualization, Writing – Original Draft. Both authors contributed to the preparation of the manuscript, read and approved the final version.

## Acknowledgements

The authors thank the researchers who generated and made the mitochondrial genome sequences publicly available in NCBI GenBank, which enabled this comparative mitogenomic and phylogenetic analysis. The authors also acknowledge colleagues who provided helpful discussions and support during the preparation of this manuscript.

## Data Availability

All sequences analyzed are publicly available in NCBI GenBank under the accession numbers in Table 1; no new sequences were generated. The pairwise distance matrix is provided as Supplementary Table S1.

## Ethical Statement

No animals were collected for this study. All analyses were performed in silico on publicly available sequence data.

